# Combined use of procalcitonin and C-reactive protein levels can help clinically diagnose bacterial co-infections in children infected with H1N1 influenza

**DOI:** 10.1101/318063

**Authors:** Zhihao Li, Liya He, Shuhua Li, Waner He, Caihui Zha, Qiaozhen Hou, Weiying Wang, Xin Sun, Huiying Liang, Wanxing Ou

## Abstract

**Objective:** This study evaluated the diagnostic value of measuring the levels of procalcitonin (PCT) and C-reactive protein (CRP) to differentiate children co-infected with H1N1 influenza and bacteria from children infected with H1N1 influenza alone and to provide a reliable clinical diagnostic support system with improved accuracy and precision control.

**Methods:** Consecutive patients (children aged <5 years) with laboratory-confirmed H1N1 influenza who were hospitalized or received outpatient care from a tertiary-care hospital in Canton, China between 1 January 2012 and 1 September 2017 were included in the present study. Laboratory results, including serum PCT and CRP levels, white blood cell (WBC) counts, and blood and sputum cultures, were analyzed. The predictive value of the combination of biomarkers versus either biomarker alone for diagnosing bacterial co-infections was evaluated using logistic regression analyses.

**Results:** Of 3180 children infected with H1N1 influenza, 226 (7.1%) met the bacterial co-infection criteria, with *Staphylococcus pneumoniae* being the most commonly identified bacteria (36.28%). Significantly higher PCT (1.46 vs 0.21 ng/ml, *p*<0.001) and CRP (19.20 vs 5.10 mg/dl, *p*<0.001) levels were detected in the bacterial co-infection group than in the H1N1 infection only group. Multivariate logistic regression analysis showed independent associations between PCT (odds ratio [OR]: 1.73, 95% confidence interval [CI],1.34-2.42, *p*<0.001) and CRP levels (OR:1.09, 95% CI, 1.06-1.13, *p*<0.001) with bacterial co-infections. Using PCT or CRP levels alone, the areas under the curves (AUCs) for predicting bacterial co-infections were 0.801 (95%CI, 0.772-0.855) and 0.762 (95%CI, 0.722-0.803), respectively. Using a combination of PCT and CRP, the logistic regression-based model, Logit(P)=-1.912+0.546 PCT+0.087 CRP, showed significantly greater accuracy (AUC: 0.893, 95%CI: 0.842-0.934) than did the other three biomarkers.

**Conclusions:** The combination of PCT and CRP levels could provide a useful method of distinguishing bacterial co-infections from an H1N1 influenza infection alone in children during the early disease phase. After further validation, the flexible model derived here could assist clinicians in decision-making processes.

## Introduction

Co-infections with bacterial pathogens are a major cause of morbidity and mortality in children with H1N1 influenza infections worldwide [1]. Most deaths that occurred during several H1N1 influenza pandemics in 1918-1919 were due to bacterial co-infections rather than direct effects of the virus[2]. A recently study estimated that bacterial co-infection was approximately 33% in patients hospitalized with H1N1infection[3], resulting in more than 74% of patients receive antibiotic therapy after admission for initial H1N1 influenza infection [4], despite adverse effects, the costs and increasing antibiotic resistance. Therefore, an early and rapid diagnosis was recognized as a priority in managing bacterial co-infections, which may assist clinicians in initiating appropriate antibiotic treatments to improve patient outcomes[5].

However, an early diagnosis of bacterial co-infections among patients with H1N1 influenza is challenging, because of the many overlapping symptoms and the lack of specific clinical manifestations of bacterial co-infections compared with H1N1 infection alone[6]; furthermore, young children cannot accurately describe their own disease symptoms, making the diagnosis even more difficult. Microbiological culture is the gold standard for diagnosing bacterial co-infections; however, current microbiological culture was time-consuming cultivation of bacteria before identification via colony and biochemical profiling, and the routine testing procedure may take several days and can also result in false-negative results.

Consequently, the availability of an efficient biomarkers system would be crucial in helping to quickly differentiate bacterial co-infections from H1N1 infections alone. Recently, several inflammatory biomarkers have been evaluated for their abilities to distinguish co-infections with H1N1 and bacteria from H1N1 infections alone. Among these biomarkers, traditional biomarkers such as a white blood cell (WBC) count [7] and C-reactive protein (CRP) levels[8] are commonly used to differentiate between bacterial and viral etiologies. Although previous studies have focused on using CRP levels to detect bacterial co-infections in patients with H1N1 infections, the evidence from these studies is inconsistent. Studies suggested serum CRP as a potential diagnostic biomarker [9–11], whereas Piacentini et al.[12] found that CRP levels were unable to distinguish bacterial co-infections from H1N1 infections. Another interesting biomarker is procalcitonin (PCT), the prohormone of calcitonin produced by C cells in the thyroid. Plasma PCT concentrations are low in healthy individuals and increase during bacterial, parasitic, or fungal infections, whereas they remain at normal levels during viral infections or noninfectious inflammatory reactions[13]. Studies have attempted to assess PCT levels in patients with H1N1 infection and found that PCT helped to distinguish bacterial co-infections from H1N1 infections[14, 15]. Nevertheless, to the best of our knowledge, previous studies published to date have focused on adults[14, 15] and patients with severe disease[16], but have included few patients with H1N1 infections.

Thus, in the present study, we conducted a retrospective analysis of 3180 children with H1N1 infection, to evaluated the diagnostic levels of serum PCT, CRP and WBC alone and in combination in differentiating bacterial co-infections from H1N1 influenza infections alone in children, to provide a reliable clinical diagnostic support system for improving diagnostic accuracy and for enabling early treatment of bacterial co-infections during H1N1 influenza infections.

## Methods

### Settings and participants

We performed a retrospective cohort study of consecutive patients with laboratory-confirmed H1N1 influenza infections, all of whom were children <5 years old who were hospitalized or received outpatient care from a tertiary-care hospital in Canton, China between 1 January 2012 and 1 September 2017. Demographic and clinical characteristics, including age, gender, weight, diagnoses, total length of hospital stay, intensive care unit (ICU) admission, total length of ICU stay, total cost and in-hospital mortality were recorded. Data from initial laboratory exams, including serum PCT and CRP levels, WBC counts, and blood and sputum cultures were collected. The ethics committee of Guangzhou Woman and Children’s Medical Center approved our study, and written informed consent was obtained from all the participants’ parents or designated guardians.

### Definitions

Patients diagnosed with H1N1 influenza infection confirmed by real-time reverse transcriptase polymerase chain reaction (RT-PCR) [17] of nasopharyngeal secretions or bronchoalveolar lavage fluid samples within the first 48 hours hospitalization were included in the study. Bacterial co-infection was defined as a positive H1N1 influenza viral PCR result with one or more bacterial pathogens detected. Bacterial cultures were obtained from blood, valid sputum, lower respiratory tract samples or samples of other normally sterile fluids within the first 48 hours hospitalization. We selected patients for this study who did not receive antibiotics prior to hospitalization to better differentiate patients co-infected with H1N1 and bacteria from patients infected with H1N1 alone.

### Inflammatory biomarkers (PCT, CRP and WBC) measurements

Venous blood samples were collected from the patients infected with H1N1 upon admission. Serum PCT levels were determined using an enzyme-linked fluorescence analysis (ELFA, VIDAS BRAHMS PCT kit, bioMerieux SA, France). Serum CRP levels were determined using BNPProSpec automatic protein analyzer (Dade Behring BN Prospec, USA)[18], and WBC counts were analyzed by using an Sysmex XE-2100 haematology analyser (Sysmex, Kobe, Japan).

### Statistical analysis

Categorical variables are summarized using absolute values and percentages, and continuous variables are presented as medians and interquartile ranges (IQR). The Chi-square tests (for nominal variables) or the Wilcoxon rank sum test (for continuous variables) was employed for between-group comparisons. Univariate logistic regression analysis was used to assess the ability of each biomarker (PCT, CRP and WBC) to diagnose bacterial co-infections. Furthermore, iterative biomarker(s) were selected (including biomarker with *p*<0.10) using automatic forwards stepwise regression, and the multivariate logistic regression model was built. The performance of the models was then assessed by calculating the area under the receiver-operating characteristic (ROC) curve(AUC). The AUC values were compared for each biomarkers individually and in conjunction with biomarkers model by Hanley and McNeil method[19]. The sensitivity, specificity, positive predictive value (PPV) and negative predictive value (NPV) were also reported. The Youden index (sensitivity + specificy-1) was used to determine the optimal ROC cutoff value. Moreover, 10-fold cross-validation to evaluate the robustness of the estimates obtained from the constructed model, as previously described [20], was performed. Then, we averaged the AUC, sensitivity and specificity values obtained from the 10-fold cross-validations to generate summary performance estimates.

All statistical analyses were performed using R Software, version 3.4.2 (www.r-project.org). A two-tailed *p* value <0.05 was considered significant.

## Results

### Study population and bacterial pathogen characteristics

During the study period, 3180 children with laboratory-confirmed H1N1 influenza infection were included, with a median age of 3.6 years (IQR, 1.8-7.5 years); 1784 (52.3%) were males. Among these patients, 226 (7.1%) had a proven bacterial co-infection. *Streptococcus pneumoniae* was the most frequent pathogens causing the bacterial co-infection in 82(36.2%) cases, followed by *Staphylococcus aureus* in 55 (24.3%) cases and *Pseudomonas aeruginosa* in 34 (15.0%) cases (Table S1). Eight children (3.5%) displayed two positive respiratory tract bacterial cultures.

When the baseline characteristics and clinical outcomes of the H1N1 plus bacterial co-infection group were compared, children in the H1N1-alone group were older, but this result was not significant (median age, 2.5 vs 2.4 years, *p*=0.197). Differences in gender or weight were not observed between the two groups; however, the bacterial co-infection group showed significantly higher inpatient admission (14.3% vs 50.4%, *p*<0.001) and ICU admission rates (2.6% vs 36.3%, *p*<0.001) than patients in the H1N1-alone group. The bacterial co-infection group also required longer hospital stays (5 vs 10 days, *p*=0.003) than H1N1-alone group and thus had much higher hospital costs (median hospital cost, 1213.2 vs 3467.3 RMB, *p*<0.001). Moreover, a higher in-hospital mortality rate was noted for the bacterial co-infection group than the H1N1 alone group (0.1% vs 4.8%, *p*<0.001) (Table 1).

**Table 1.**
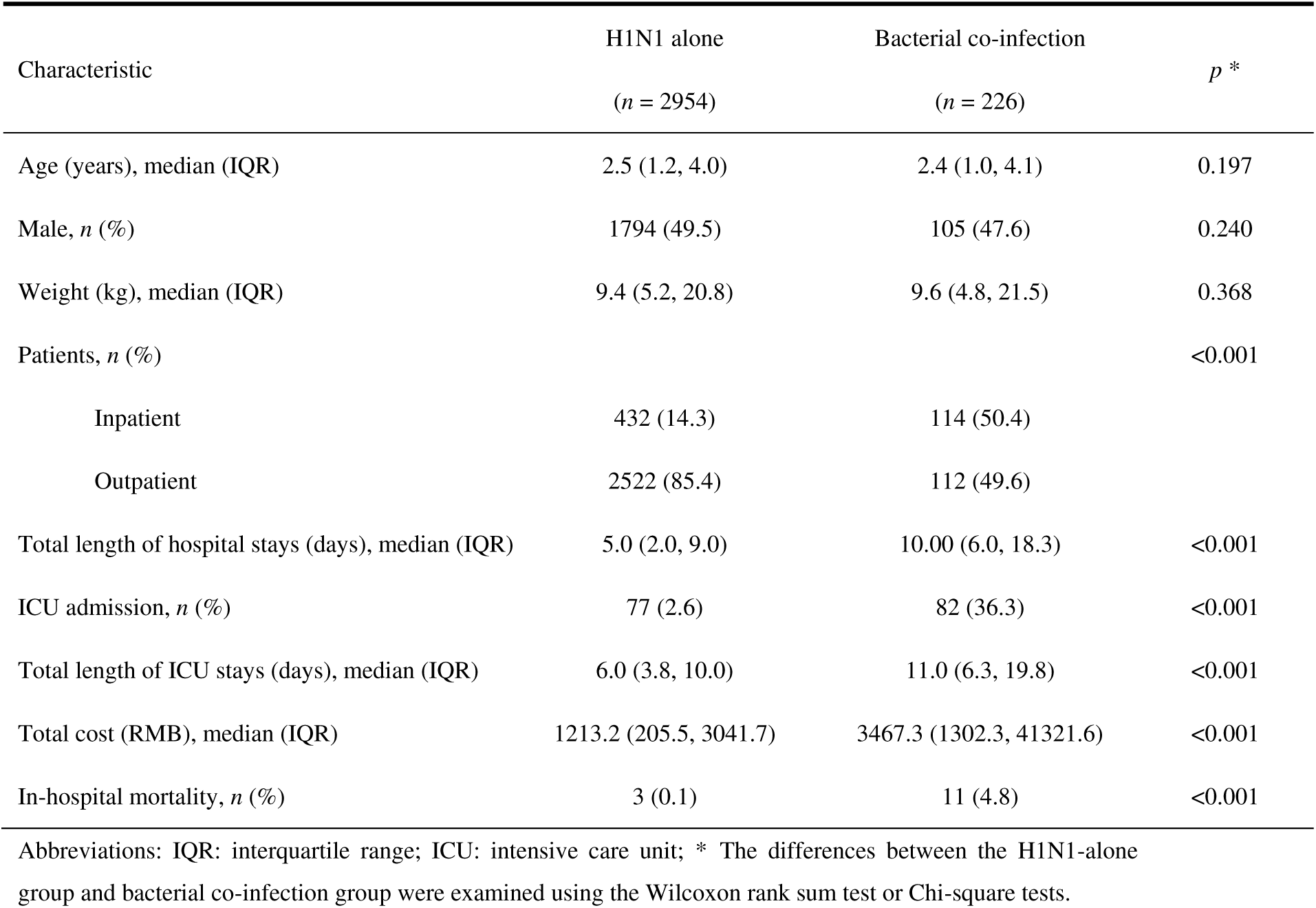
Demographic and clinical characteristics of patients with H1N1 influenza who presented with and without bacterial co-infections.

### Comparison of serum PCT, CRP and WBC levels between H1N1-alone and H1N1 with bacterial co-infection groups

Serum PCT, CRP and WBC levels were analyzed to identify potential biomarkers that distinguished between H1N1 infections and H1N1 and bacterial co-infections. The median serum PCT, CRP and WBC levels were all significantly higher in the H1N1 with bacterial co-infection group than in the H1N1-alone group (median PCT level, 1.46 vs 0.21 ng/ml, *p*<0.001; median CRP level, 19.20 vs 5.10 mg/dl, *p*<0.001, median WBC count, 8.50 vs 6.90 ×10^9^ cells/l, *p*=0.019) (Figure 1).

**Figure 1.**
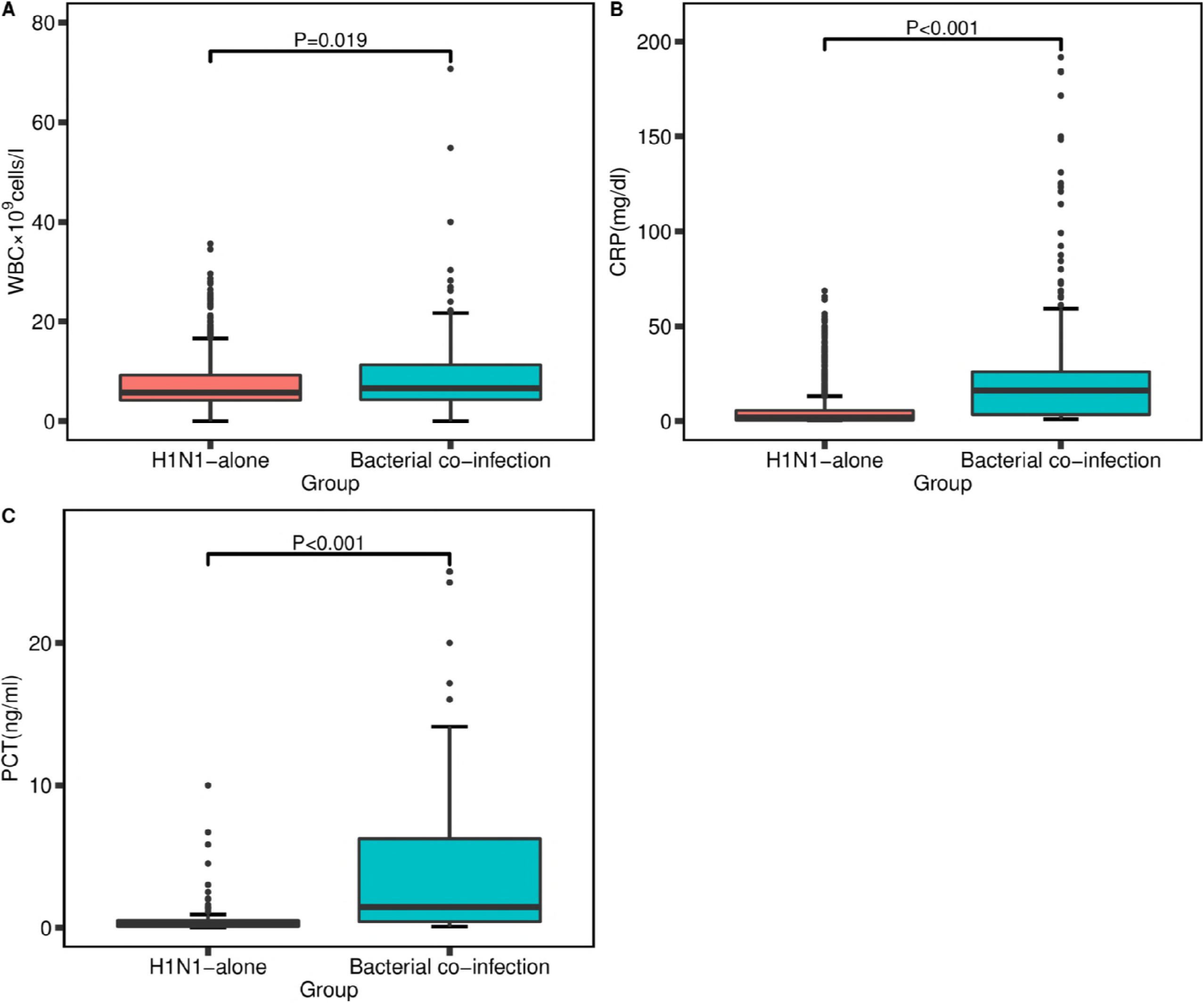
Serum PCT (A), CRP (B) and WBC (C) levels in patients with H1N1 influenza who presented with and without bacterial co-infections. The differences between the H1N1-alone group and bacterial co-infection group were examined using the Wilcoxon rank sum test.

### Univariate and multivariate logistic regression analyses

Univariate analysis revealed significant associations of serum PCT, CRP and WBC levels with co-infections with H1N1 and bacteria (odds ratio [OR]:1.65, 95 % confidence interval [CI] 1.34-2.06, *p*<0.001; OR: 1.08, 95 % CI 1.06-1.09, *p* < 0.001; OR:1.06, 95% CI 1.04-1.09, *p*=0.02, respectively). The associations with PCT and CRP levels remained statistically significant(*p*<0.05) after the application of the forwards regression model, whereas WBC counts were excluded from the model (*p* <0.05). Then, multivariate logistic regression analysis showed that CRP (OR:1.09, 95% CI 1.06-1.13, *p*<0.001) and PCT levels (OR:1.73, 95%CI 1.34-2.42, *p*<0.001) were significant independent diagnostic biomarkers. (Table 2).

**Table 2.** Univariate and multivariate logistic regression analyses of biomarkers for of bacterial co-infection in H1N1 patients infected with H1N1.

| Variable | Univariate |  |  |  | Multivariate |  |  |  |
| --- | --- | --- | --- | --- | --- | --- | --- | --- |
| | $\beta$ | OR | 95% CI | *p* | $\beta$ | OR | 95% CI | *p* |
| WBC | 0.060 | 1.06 | 1.04-1.09 | <0.001 | -* | - | - | - |
| PCT | 0.498 | 1.65 | 1.34-2.06 | <0.001 | 0.546 | 1.73 | 1.34-2.42 | <0.001 |
| CRP | 0.073 | 1.08 | 1.06-1.09 | <0.001 | 0.087 | 1.09 | 1.06-1.13 | <0.001 |

Abbreviations: $\beta$ : regression coefficient; OR: odds ratio; CI: confidence interval; \* In the multivariate logistic regression analysis, WBC counts ( $p>0.05$ ) were excluded from the final model based on the results of the forward stepwise analysis.

### Comparison and validation of the model’s diagnostic ability

Because the serum PCT and CRP levels were independent predictors that differentiated patients with bacterial co-infections from patients infected with H1N1 alone, we constructed a new model, PCT&CRP [Logit (P) = -1.912 + 0.546 PCT+ 0.087 CRP], that combined the PCT and CRP levels. The performance of the ROC curves of the constructed model, PCT, CRP, and WBC levels for differentiating children with H1N1 and bacterial co-infections from children infected with H1N1 alone were compared. The AUC, sensitivity, specificity, PPV, and NPV are shown in Table 3. The constructed model exhibited the largest AUC (0.893, 95%CI 0.852-0.934). The *p* values of the ROC curve comparison between the constructed model and CRP and PCT levels were all less than 0.01. The AUCs for PCT, CRP, and WBC levels were 0.801(95%CI, 0.772-0.855), 0.762(95%CI, 0.722-0.803), and 0.551(95%CI, 0.502-0.592), respectively. The optimum cutoff values for PCT, CRP, and WBC were 0.52 ng/ml, 13.55 mg/l and 11.56×10^9^ cells/l, respectively. Significant differences were observed among the ROC curves of the PCT, CRP, and WBC (*p*<0.05). The diagnostic ability of each model followed the order of PCT&CRP > PCT > CRP > WBC (Figure 2). The PCT&CRP was superior to use of the PCT, CRP and WBC alone in differentiating patients with bacterial co-infections from those infected with H1N1 alone. The robustness of PCT&CRP was internally evaluated through 10-fold cross-validation. On average, the constructed model presented an AUC of 0.872, a sensitivity of 0.754 and a specificity of 0.896.

**Table 3.** Discriminatory performance of WBC, CRP, PCT and the constructed model for detecting patients with H1N1 influenza and a bacterial co-infection.

| Variables | AUC (95% CI) | cutoff level | Sensitivity | Specificity | PPV | NPV |
| --- | --- | --- | --- | --- | --- | --- |
| WBC | 0.551 (0.502-0.592) | 11.56 | 0.267 | 0.887 | 0.144 | 0.910 |
| CRP | 0.762 (0.722-0.803) | 13.55 | 0.633 | 0.856 | 0.330 | 0.971 |
| PCT | 0.801 (0.772-0.855) | 0.52 | 0.643 | 0.886 | 0.773 | 0.852 |
| PCT&CRP * | 0.893 (0.852-0.934) | - | 0.830 | 0.868 | 0.854 | 0.846 |

Abbreviations: AUC: area under the receiver-operating-characteristic curve; CI: confidence interval; PPV: positive predictive value; NPV: negative predictive value; PCT&CRP: Logit (P) = $-1.912 + 0.546 \text{ PCT} + 0.087 \text{ CRP}$ .

**Figure 2.**
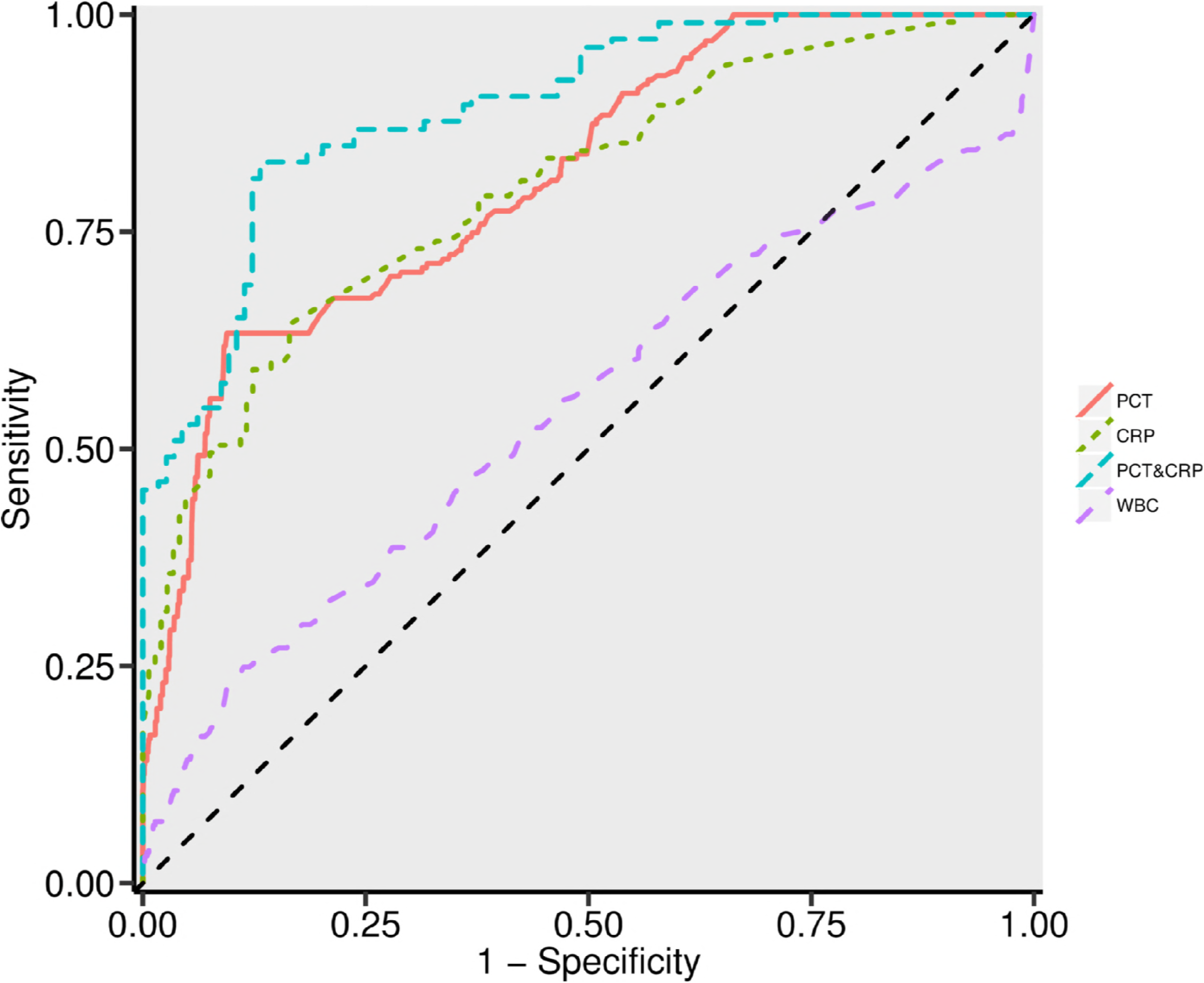
ROC curves of PCT, CRP, WBC and PCT&CRP (Logit (P) = -1.912 + 0.546 PCT+ 0.087 CRP) for differentiating patients with bacterial co-infections from those with infected H1N1 alone.

## Discussion

Bacterial co-infection is especially known to excess the mortality and morbidity of H1N1 influenza. Unfortunately, it is difficult to correctly diagnose bacterial coinfection based only on clinical criteria, as well as bacterial culturing is time-consuming. There is a crucial need to differentiate H1N1 patients with bacterial co-infection from H1N1 infection alone. The diagnostic and predictive value of serum PCT and CRP levels as biomarkers has been discussed in several studies[10, 14, 15, 21]. Shin et al. [10]found that serum PCT was a good indicator of in discriminating bacterial co-infections infection from H1N1 infections alone in 60 adult patients in ICU. Guervilly et al. [21] report that PCT values were statistically higher in patients with bacterial co-infections. In addition, PCT has been suggested to exclude bacterial co-infections in patients with an H1N1 infection and to reliably and accurately reduce inappropriate antibiotic exposure [14]. Our results showed that serum PCT levels was significantly higher in patients with bacterial co-infection compared to those infected H1N1 infection alone, reminding us that PCT was association with bacterial co-infection. Furthermore, the results of ROC curve analysis indicated that an AUC value of 0.801 (95% CI, 0.772-0.855), with cutoff value was 0.52ng/ml, supported the prognostic value of PCT in children with or with bacterial co-infection.

The diagnostic utility of CRP to differentiate bacterial co-infection from H1N1 infection is disputed [9–11, 15]. Haran et al. [11]found that CRP as predictor of bacterial infection among patients with an H1N1 infection. Similarly, Shin et al. [10] reported that serum CRP levels was significantly higher in patients with bacterial co-infection compared to those infected with H1N1 alone. But other study suggested that CRP levels were unable to distinguish bacterial co-infections from H1N1 infections [9]. Our study showed that serum CRP levels was significantly higher in patients with bacterial co-infection compared to those infected H1N1 infection alone, it indicating that those biomarkers could be aid in discriminating between these conditions. Furthermore, our study showed that the diagnostic efficacy of PCT for bacterial co-infection in H1N1 infection was better than that of CRP (AUC 0.801 and 0.783, respectively; p<0.05), consistent with the results of a previous study[11]. However, the AUC of WBC counts in diagnosing bacterial co-infections was 0.551 (95%CI, 0.502-0.592), indicating that WBC may not be a valuable biomarker for our cohort of children.

Previous study use of a combination of CRP and PCT levels for evaluating for bacterial co-infections increased the accuracy of differentiating children with bacterial co-infections from those infected with H1N1 alone[10]. Similar observations were reported in the present study, we used a multivariate logistic regression analysis to construct a new model using the PCT and CRP levels: [Logit(P)=-1.912+0.546 PCT+0.087 CRP]. The ROC curve analysis yielded an AUC value for the model of up to 0.893, which was clearly superior to PCT or CRP levels alone (p<0.05). Furthermore, the constructed model [Logit(P) =-1.912+0.546PCT +0.087CRP] was internally validated through 10-fold cross-validation, resulting in high diagnostic accuracy. Therefore, the joint detection of PCT and CRP levels clearly improves the prognosis of children with H1N1 bacterial co-infection. Based on the results from our study, the combination of serum PCT and CRP levels will help clinicians determine the appropriate antibiotic therapy[22], thus potentially improving patient outcomes and reducing antibiotic overuse [5].

This study involved 3180 children with H1N1, 7.11% of whom presented a confirmed bacterial co-infection, after including both outpatients and inpatients. The proportion of bacterial co-infection while similar to that previously reported for H1N1. Nevertheless, previous studies of children with H1N1 influenza infection reported a bacterial co-infection rate ranging from 18% to 60%[23, 24]. These rates may be overestimated because the previous studies were limited to pediatric patients in the ICU, which represent a population with moderate to severe H1N1 influenza infection. Moreover, children with bacterial co-infections exhibited a higher percentage of ICU admission rates in the current study.

Our study shows that *Streptococcus pneumoniae* was the leading cause of bacterial co-infection with H1N1, followed by *Staphylococcus aureus* and *Streptococcus pyogenes,* consistent with the results from previous studies[25, 26]. Additionally, children with H1N1 influenza infection and bacterial co-infection have been reported to exhibit a higher risk of severe outcomes [26–28]. In our study, patients co-infected with bacteria and H1N1 exhibited increased percentages of inpatient and ICU admissions, higher costs and longer hospital stays. Furthermore, a significantly higher hospital mortality rate was observed in children with H1N1 and bacterial co-infections because bacterial co-infections represent an important mortality risk factor, possibly suggesting early empiric antibiotics treatment in severe patients may improve outcomes.

The potential limitations of our study should be mentioned. First, the levels of selected biomarkers (PCT, CRP and WBC) were evaluated only once. Second, our diagnostic model was derived and validated at a single hospital center, and it should be validated in a multicenter trial center before its broad application. Finally, we also acknowledge that, we may have created a bias, due to bacterial organisms cannot be confirmed solely with blood, sputum, lower respiratory tract samples.

## Conclusion

In conclusion, we detected serum PCT and CRP levels and revealed that they represent promising biomarkers and useful clinical tools for differentiating pediatric patients with bacterial co-infections from those infected with H1N1 alone. Furthermore, the combination of PCT and CRP levels could represent a useful method for screening bacterial co-infections from H1N1 influenza infections alone in children during the early disease phase. After further validation, the flexible model reported here may assist clinicians with decision-making processes.

## Acknowledge

This research did not receive any specific grant from funding agencies in the public, commercial, or not-for-profit sectors.

## Supplementary material

**Table S1** Pathogens isolated in patients with H1N1 influenza and a bacterial co-infection

